# A living organism-level model of subcellular architecture across adult life reveals a midlife architectural state shift

**DOI:** 10.64898/2026.11.11.765956

**Authors:** Christopher S. Morrow, Aditi Prabhakar, Jamie D. Kraft, Carlos Giovanni Silva-García, Praju Vikas Anekal, Paula Montero Llopis, Jeffrey W. Miller, William B. Mair

## Abstract

Many features of subcellular organelle architecture change throughout life and influence aging. However, our understanding of this structural dynamism relies on single cell systems or comparisons between data sets focused on a limited number of organelles or timepoints. To overcome this barrier, we developed Multiorganelle Mapper (**MoMA**), a *C. elegans* strain enabling simultaneous imaging of six organelles across all somatic cell types in fully intact, living animals via fluorescence microscopy. We use MoMA to generate an organism-level *in vivo* map of subcellular architecture throughout adult life to understand how and why organelle organization changes with age. Aging drives a widespread and consistent shift in subcellular architecture throughout the body. Strikingly, most of this remodeling does not accumulate progressively, but emerges abruptly within individuals at midlife, an inflection point we term the midlife architectural state shift (**MASS**). Through a targeted RNAi screen, we show the MASS is not unique to aging, but instead resembles a generalized reactive state which can be triggered by a range of distinct homeostatic challenges. Finally, we show the MASS both coincides with and can be driven by midlife physiological changes, suggesting that this remodeling may largely reflect a coordinated reaction to the aged organismal state. Together, we present MoMA as a practical approach for visualizing subcellular architecture at organismal scale and demonstrate how it can be used to uncover principles underlying the dynamic regulation of subcellular architecture throughout life.

## Introduction

Many biological processes are tightly coupled to the dynamically regulated abundance, morphology and organization of organelles distributed throughout the cytoplasm (termed subcellular architecture)^1–12^. For example, endoplasmic reticulum (**ER**) morphology in the liver governs metabolic homeostasis^13^, while contact sites between peroxisomes and mitochondria support mitochondrial redox homeostasis^14^. Moreover, numerous features of subcellular architecture change throughout life and can be targeted to modulate longevity^15–23^. For instance, many longevity interventions have been found to extend lifespan through tuning the architectural properties of mitochondria^15,19^ and nucleoli^16^. Yet our understanding of this structural plasticity remains constrained by the tools available to visualize it. For example, imaging many organelles simultaneously in living samples currently requires studying cells *in vitro*^24,25^. Although informative, *in vitro* systems isolate the cell from the rest of the body, removing extrinsic cues that can substantially alter cellular behavior. Mitochondria, for instance, often fuse in response to fasting *in vitro*^26^ but were recently shown to fragment in response to fasting *in vivo*^27^. Fixed-tissue approaches such as electron microscopy, by contrast, can resolve many organelles at high resolution *in vivo* but cannot capture organelle dynamics in living cells, limiting the questions they can address^13,28^. Moreover, because it remains practically challenging to analyze many cell types in a single experiment, most studies to date have focused on one cell type or tissue and do not capture the diversity of cell types and tissues which exist across the body of an organism. As a result, it remains unclear whether age-dependent changes reported in specific cell types are restricted to those cells or shared throughout the body, and how these changes relate to one another in space and time across organelles, cell types and tissues. Ultimately, these gaps together heavily limit our ability to determine both why subcellular architecture changes with age and what those changes mean for the animal.

To overcome these barriers, we developed MoMA, a single *C. elegans* strain that enables simultaneous visualization of six organelles across somatic cell types in fully intact, living animals. Each organelle is labeled with a distinct fluorescent protein expressed via genomically integrated transgenes, so subcellular architecture can be imaged by fluorescence microscopy immediately after immobilization, without time-consuming or costly sample preparation, making this a highly accessible approach. Here we use MoMA to map age-dependent changes in organelle morphology and abundance across the body, and use these data to better understand how and why subcellular architecture remodels during aging. Unlike other features of cell biology that develop progressively and stochastically with age^29–33^, we find that most age-associated remodeling of subcellular architecture emerges relatively abruptly and predictably at an inflection point in midlife, in an event we term the MASS. Further, we show that much of this aged state is not unique to aging, but instead is broadly inducible by a range of homeostatic challenges and may reflect a reaction to a midlife shift in physiology. Together, here we establish MoMA as an accessible, organism-level platform for visualizing subcellular architecture in living animals and demonstrate how it can reveal principles governing the dynamic regulation of subcellular architecture across life. Because MoMA resolves many layers of cytoplasmic organization in their native tissue context across an intact animal, and can be applied to virtually any physiological condition, we anticipate that MoMA and the datasets generated here will be valuable resources for decoding the spatial logic of cell biology in aging, health and disease, and for training artificial intelligence models toward predictive, *in silico* representations of the *in vivo* cellular landscape and its response to age and perturbations.

## Results

### MoMA enables visualization of 6 organelles simultaneously across somatic cell types in fully intact living *C. elegans*

To develop a model to easily, rapidly and simultaneously visualize subcellular architecture at the organismal level within intact, living animals, we assembled transgenic *C. elegans* strains harboring widely expressed transgenes composed of spectrally distinct fluorescent proteins attached to organelle-targeting peptides, similar to systems used in cell lines *in vitro*^25,34,35^. Specifically, we created a single strain harboring: 1) wrmBFP::PLIN-1; labeling lipid droplets (endogenous tag)^36^, 2) HIS-24::eCFP; labeling nuclei (integrated extrachromosomal array driven by *eft-3p*)^37^, 3) TOMM-20^aa^^1–49^::GFP; labeling mitochondria (single copy knock-in driven by *eft-3p*)^38^, 4) TRAP-1::hfYFP; labeling the endoplasmic reticulum (integrated extrachromosomal array driven by *eft-3p*)^39^, 5) CemOrange2::SKL; labeling peroxisomes (integrated extrachromosomal array driven by *eft-3p*)^21,40^ and 6) NUC-1::mCherry, labeling lysosomes (integrated extrachromosomal array driven by *eft-3p*)^41^, which we collectively call MoMA (Fig. 1A-C; more details on MoMA are provided in the methods). Each reporter was widely expressed across the body, and we could reliably discriminate all labels using two distinct confocal imaging strategies: either by precisely controlling excitation and emission wavelengths (Fig. S1A–C) or by using spectral imaging paired with linear unmixing (Fig. S1D–H). Comparing each MoMA reporter to established markers of the corresponding organelle, we confirmed that each label localized to the intended compartment and did not overtly perturb organelle morphology (Fig. S1I–J). Thus, MoMA provides a stable system for visualizing subcellular architecture across the *C. elegans* body at organismal scale in living animals, without additional sample processing or costs beyond standard animal husbandry.

**Figure 1.**
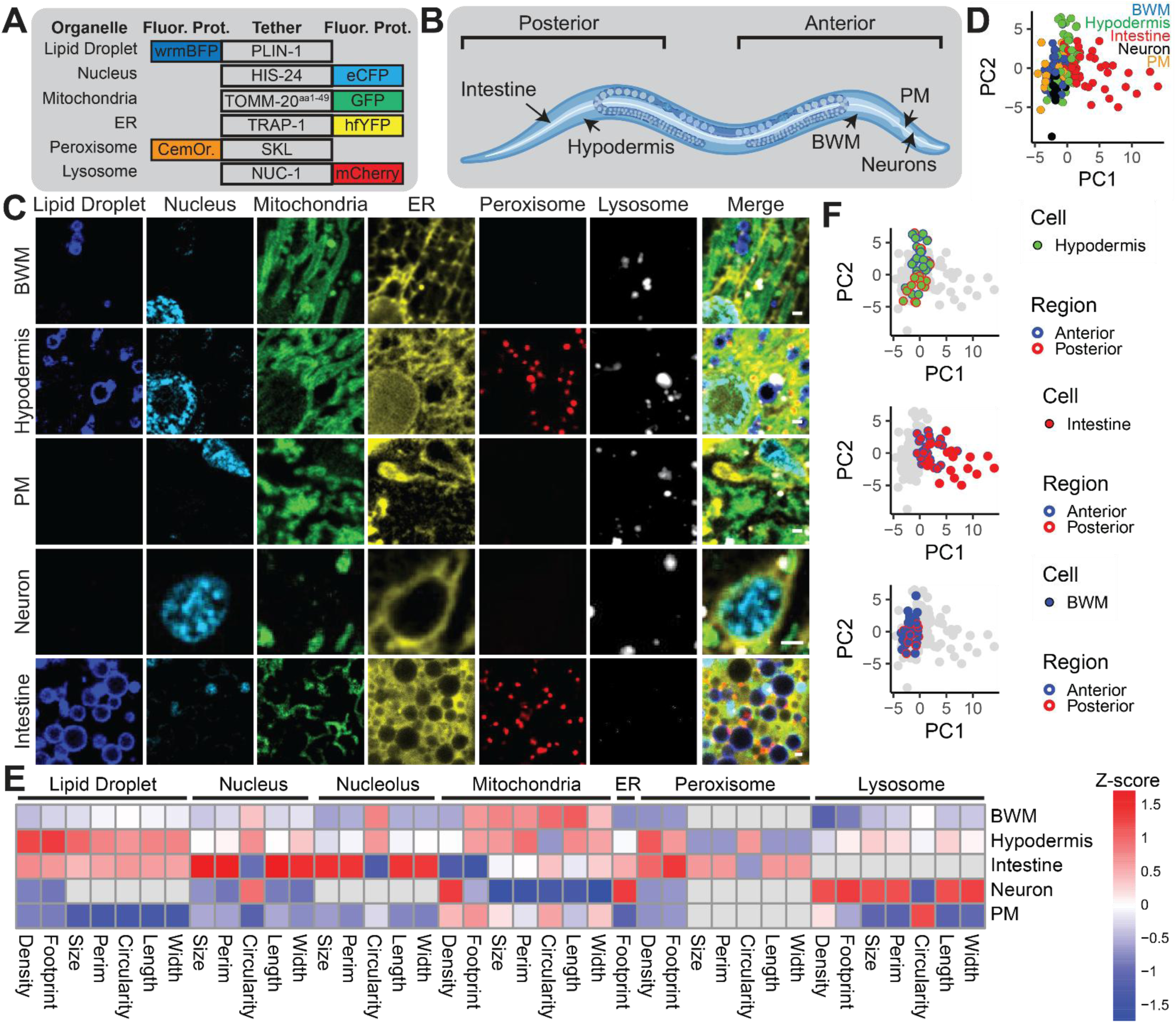
Multiorganelle Mapper (MoMA): A living, animal-scale model of subcellular architecture. A) Schematic listing transgenes composed of spectrally distinct fluorescent proteins fused to organelle tethering peptides in MoMA *C. elegans* used to label 6 organelles across somatic cell types simultaneously. CemOrange2 is abbreviated as CemOr. B) Schematic showing cell types and body regions throughout the *C. elegans* body which were imaged in this study. Created in BioRender. Morrow, C. (2024) https://BioRender.com/v72k420. C) Images of lipid droplets (wrmBFP::PLIN-1), nuclei (HIS-24::eCFP), mitochondria (TOMM-20_aa1-49_::GFP), ER (TRAP-1::hfYFP), peroxisomes (CemOrange2::SKL) and lysosomes (NUC-1::mCherry) across neuronal, BWM, hypodermal, intestinal and PM cells in day 2 adult *C. elegans*. D) Plot of PC1 versus PC2 from a PCA of all architectural endpoints per cell across anterior BWM, hypodermal, intestinal, neuronal and PM cells in day 2 adult *C. elegans*. Individual dots represent individual cells (n=27 animals, 23-27 cells per cell type). E) Heat map showing Z-score of individual architectural endpoints across organelle type and feature (columns) and anterior cell types (BWM, hypodermis, intestinal and neuronal cells; rows), normalized across cell types in day 2 adult *C. elegans* (n=27 animals, 23-27 cells per cell type). F) Plot of PC1 versus PC2 from a PCA of all architectural endpoints per cell across neurons, PM cells and anterior and posterior BWM, intestinal and hypodermal cells in day 2 adult *C. elegans*. Subsets of cell types are labeled by cell identity and body region in each plot respectively. Each dot represents an individual cell (n=27 animals, 23-27 cells per cell type). Scale bars, 1 µm.

### Aging drives widespread changes in subcellular architecture across the body which develop largely consistently across individuals

We first used MoMA to generate a data set enabling us to map age-dependent changes in subcellular architecture across the body. We imaged major somatic cell types in day 2, 6, 9, 12 and 15 old MoMA *C. elegans,* including: neurons, pharyngeal muscle (**PM**) cells, hypodermal cells, intestinal cells and body wall muscle (**BWM**) cells in multiple regions of the body (Fig. 1B-C; cell type validation in Fig. S1K). This data set is particularly powerful, both because of the breadth of architectural features it captures and because every measurement stays tied to its location in the body, with each organelle indexed to the cell type, body region and individual animal in which it was imaged, so that any feature can be compared against any other elsewhere in the same animal. To our knowledge, no other data set analyzing aging animals retains this level of spatial context across the body of an animal. To begin exploring these data, we used an analytic pipeline focused on organelle geometry and abundance to quantify differences in subcellular architecture across organelle types, cell types, body regions, individuals and ages (Fig. 1B-F, S1L).

We first asked how much subcellular architecture varied across cells throughout the body. We found that subcellular architecture was in many respects highly cell-type specific, with organelles having a variety of variable morphological features across somatic cell types and some cell types looking more similar than others (Fig. 1B-E, S1L). Lipid droplets and peroxisomes were primarily limited to hypodermal and intestinal cells, whereas mitochondria, nuclei, ER and lysosomes appeared more ubiquitously across cell types, albeit with varying morphological features. Analyzing cells imaged in either anterior or posterior regions of the animal, we found that cells of each type generally clustered together, suggesting that subcellular architecture is largely conserved within cell types across the body (Fig. 1F). Together, our data show that subcellular architecture is strongly cell-type specific and, despite some exceptions, generally conserved within cell types across regions throughout the body.

We next asked how subcellular architecture changes across different cell-types and body regions during aging by comparing measurements made in day 2 and day 12 animals. We found that aging induced significant changes in features of subcellular architecture spanning many organelles and cell types (Fig. 2A-B, S2A-C). Analyzing anterior hypodermal cells for example, we found that on average lipid droplets became more abundant, mitochondria fragmented, ER footprint decreased and lysosome size increased. Interestingly, despite the fact that subcellular architecture is distinct across cell types, we found that many of these changes in organelle morphology and abundance were consistent across cell types and body regions. For instance, though the amplitude was variable, most cell types exhibited fragmentation of mitochondria with age, reduction in ER footprint and expansion of the lysosome pool (Fig. 2B-D). However, highlighting the importance of analyzing subcellular architecture across cell types and body regions using a system such as MoMA, other organelles showed cell-type and body-region specific changes with age. For instance, lipid droplets accumulate strongly in anterior hypodermal cells, while becoming depleted in anterior intestinal cells and remaining largely unchanged in posterior hypodermal cells (Fig. 2E-F, S2A, D-E). Most changes we resolved were consistent with expectations based on prior work^15–22^. However, some observations were surprising. For example, smaller nucleoli have been associated with longevity^16^, and nucleolar enlargement has been reported in multiple age-related contexts^22^, but we observed no significant change in nucleolar size between young and old animals in any cell type (Fig. 2B). Importantly, architectural changes detected by MoMA were consistently validated using independent fluorescent reporters, supporting the accuracy of MoMA in capturing age-dependent subcellular remodeling and indicating that these discrepancies likely reflect biological differences rather than technical artifacts linked to MoMA (Fig. S2F). In sum, these data show that aging drives widespread changes to subcellular architecture across the body, some of which are context specific, but many of which are generalized across cells throughout the body. More broadly, our data suggest that there may be many generalized architectural strategies that cells employ to react to the challenges imposed by aging.

**Figure 2.**
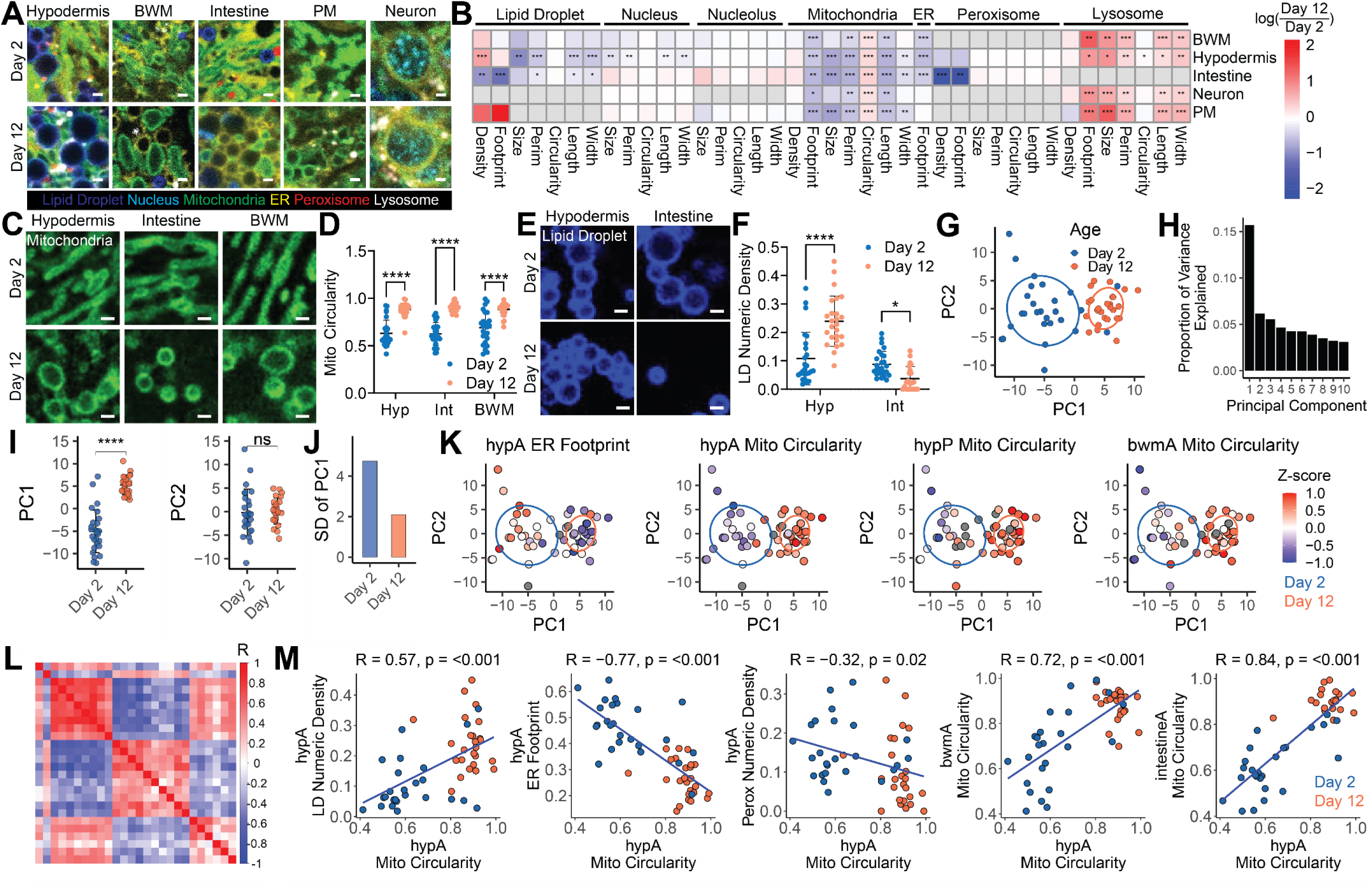
Aging drives a widespread and consistent shift in subcellular architecture throughout the body. A) Images of day 2 or day 12 adult MoMA *C. elegans* hypodermal, BWM, intestinal, PM and neuronal cells. B) Heat map showing log(day 12/day 2) of architectural endpoints analyzed across day 2 and day 12 old MoMA *C. elegans* anterior BWM, hypodermal, intestinal, neuronal and PM cells (n=26-27 animals per age, moderated t-test, mean ± SD). C-D) Images and analysis of mitochondrial circularity in day 2 or day 12 MoMA *C. elegans* hypodermal, intestinal and BWM cells (n=22-26 cells per group; Two-way ANOVA with post-hoc Tukey’s test, mean ± SD). E-F) Images and analysis of lipid droplet density in day 2 or day 12 MoMA *C. elegans* hypodermal and intestinal cells (n=22-26 cells per group, Two-way ANOVA with post-hoc Tukey’s test, mean ± SD). G) Plot of PC1 versus PC2 from a PCA of all 312 architectural endpoints analyzed per animal across day 2 and day 12 adult MoMA *C. elegans* (n=26-27 animals per age; PCA). H) Plot of proportion of variance attributed to each of the first ten principal components from the PCA shown in Fig. 2G. I) Plots of day 2 and day 12 MoMA *C. elegans* PC1 and PC2 values (n=26-27 animals per age, Student’s t-test, mean ± SD). J) Plot showing standard deviation across PC1 from the PCA shown in Fig. 2G across either day 2 or day 12 MoMA *C. elegans* (n=26-27 animals per age). K) Plots of the PCA shown in Fig. 2G with individuals colored by features of subcellular architecture (n=26-27 animals per age, PCA). L) Correlation matrix showing the correlation between a subset of architectural endpoints that change significantly with age across organelle types, cell types and body regions in day 2 and day 12 MoMA *C. elegans* (n=26-27 animals per age, Pearson Correlation). M) Correlation plots showing architectural endpoints relative to each other across individuals in day 2 and day 12 MoMA *C. elegans*. Each point displays an individual *C. elegans* (n=26-27 animals per age, Pearson Correlation). Scale bars, 1 µm. ∗ p < 0.05, ∗∗ p < 0.01, ∗∗∗ p < 0.001, ∗∗∗∗ p < 0.0001, for heatmaps, significance is capped at three asterisks.

Aging is often thought to be a highly heterogeneous process, because of the observation that many biological measurements become progressively more heterogeneous during aging^30–33^. Therefore, we were curious to understand whether subcellular architecture would also become more heterogeneous during aging, or whether these changes may be more coordinated and predictable. We performed a principal component analysis (**PCA**) of all features of subcellular architecture recorded per animal across day 2 and day 12 animals. We found that day 12 animals exhibited a strong shift along a single principal component in which they were almost entirely separated from day 2 animals (Fig. 2G-I). Surprisingly, we found that aged animals harbored subcellular architectures that were, if anything, more homogenous across PC1 and PC2 compared to young animals (Fig. 2J). This observation suggests that many features of subcellular architecture covary together, predictably, as opposed to developing independently. Analyzing individual features across animals, we also observed similar patterns. Though subcellular architecture is not always strictly synchronized in either a youthful or aged-like state, many feature changes appeared relatively consistently across and within cells throughout day 12 individuals (Fig. 2K-M). Thus, in contrast with many other types of biological features that become progressively more heterogeneous during aging^30–32^, our data together suggest that aging induces many architectural changes which are generally consistent across individuals.

### Age-dependent changes to subcellular architecture precipitate abruptly in midlife within individuals but with stochastic timing across individuals

We next asked when features of subcellular architecture change across the body during aging, by analyzing day 2, 6, 9, 12 and 15 animals relative to each other. Interestingly, initial qualitative inspection suggested that across all ages animals largely exist in one of two predominant architectural states: a youthful-like or aged-like configuration. Consistent with this, principal component analysis revealed that most day 6 animals closely resembled day 2 animals, while day 12 and day 15 animals were highly similar to each other (Fig. 3A-C). In contrast, day 9 animals were heterogeneous, with some clustering alongside youthful animals, some alongside older aged animals, and a minority of others that appeared to be transitioning from a youthful to an aged-like state. This pattern suggests that many architectural features generally do not change gradually over time, but instead shift relatively abruptly within a restricted window in midlife. Analysis of individual features further supported this interpretation. Multiple endpoints, including anterior hypodermal lipid droplet density, mitochondrial circularity, and ER footprint, remained relatively stable between day 2 and day 6, and between day 12 and day 15, but exhibited pronounced changes in midlife, and in some cases even appeared loosely bimodally distributed across many Day 9 old individuals (Fig. 3D-E, S3A-B). Together, these data indicate that the widespread architectural remodeling we resolve occurs over a relatively acute period in midlife rather than progressively throughout life.

**Figure 3.**
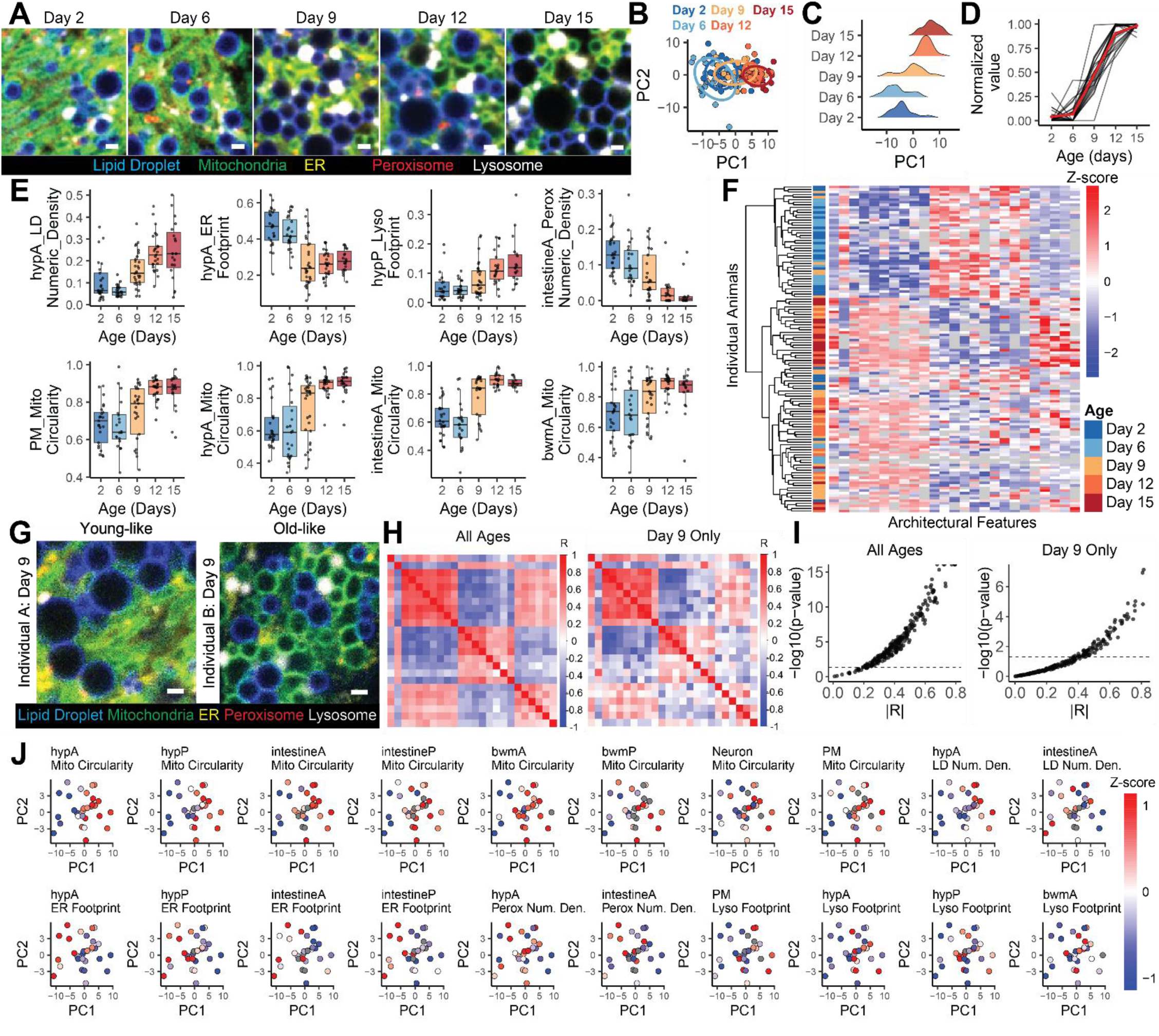
Age-dependent changes to subcellular architecture develop relatively abruptly at an inflection point in midlife within individuals as a part of the “midlife architectural state shift (MASS)”. A) Images of day 2, 6, 9, 12 and 15 adult MoMA *C. elegans* anterior hypodermal cells. B) Plot of PC1 versus PC2 from a PCA of all 312 architectural endpoints analyzed per animal across day 2, 6, 9, 12 and 15 adult MoMA *C. elegans* (n=128 animals, 20-30 animals per age, PCA). C) Histogram of PC1 across day 2, 6, 9, 12 and 15 animals from the PCA in Fig. 3B (n=128 animals, 20-30 animals per age, PCA). D) Plots of features of subcellular architecture which significantly change during aging across day 2, 6, 9, 12 and 15 animals with youthful values set to 0 and aged values set to 1. A red line shows the average of all feature changes presented in this plot (n=128 animals, 20-30 animals per age). E) Plots of individual features of subcellular architecture across day 2, 6, 9, 12 and 15 MoMA *C. elegans* (n=128 animals, 20-30 animals per age, box shows median and interquartile range). F) Heatmap showing features of subcellular architecture which significantly change during aging (down columns) across individual animals (across rows), with age annotated across day 2, 6, 9, 12 and 15 MoMA *C. elegans* (n=128 animals, 20-30 animals per age). G) Images of anterior hypodermal cells across two distinct MoMA *C. elegans* which are either more reminiscent of youthful or aged-like animals. H) Correlation matrices analyzing the correlation between features of subcellular architecture that change significantly during aging across either all ages of animals analyzed or just day 9 animals (n=128 animals, Pearson Correlation). I) Plot of the absolute value of Pearson’s correlation (r) compared against the -log10(p-value) reflecting significance of correlation across either all ages of animals analyzed or just day 9 animals (n=128 animals, Pearson Correlation). J) Plots of PC1 vs PC2 from the PCA shown in Fig. 3B showing day 9 animals only, colored by values of individual features of subcellular architecture (n=30 animals, PCA). Scale bars, 1 µm.

We next asked whether these distinct features of aged subcellular architecture develop independently or in a coordinated manner within individuals and whether certain changes may consistently precede or follow others. As previously noted, within day 9 animals, many individuals qualitatively appeared globally youthful-like or aged-like across multiple cell and organelle types, rather than exhibiting random mixtures of features. Consistent with this observation, when examined quantitatively across individuals, features that changed with age tended to covary, even across distinct organelles, tissues, and body regions (Fig. 3F-J). This coordination was evident both when analyzing all ages together and, importantly, even just within day 9 animals alone, where numerous feature pairs remained significantly correlated. Interestingly, though these distinct events did not emerge strictly simultaneously, we did not observe any clear examples of specific organelles and/or cell types consistently changing before or after others. Thus, while animals undergo this structural reorganization relatively abruptly in midlife, the order in which changes manifest across the body can still vary between individuals. Together, these findings indicate that age-dependent remodeling of subcellular architecture is not purely stochastic, but instead likely reflects an organism-wide transition triggered by a systemically experienced midlife event; we term this transition the midlife architectural state shift (**MASS**).

### A targeted RNAi screen reveals that many features of the MASS are broadly inducible responses to disruption in homeostatic mechanisms and are frequently coregulated within individuals, as seen during aging

The relative temporal synchrony in which MASS features develop within individuals implies that there may be a shared upstream driver of many of these changes, yet what this driver may be remains unclear. Therefore, to gain insight into which cellular processes may causally regulate MASS features, we next profiled subcellular architecture in young adults exposed to a panel of RNAi conditions targeting a wide range of distinct cellular processes throughout various cellular compartments – lipid synthesis (*fasn-1*), nucleolar regulation (*ncl-1*), electron transport chain (*cco-1*), peroxisome function (*prx-5*), energy sensing (*aak-2*/AMPK), tricarboxylic acid cycle (**TCA**) cycle (*aco-2*), and glycogen synthesis (*gsy-1*) – and compared these architectures to those observed during aging (Fig. 4).

**Figure 4.**
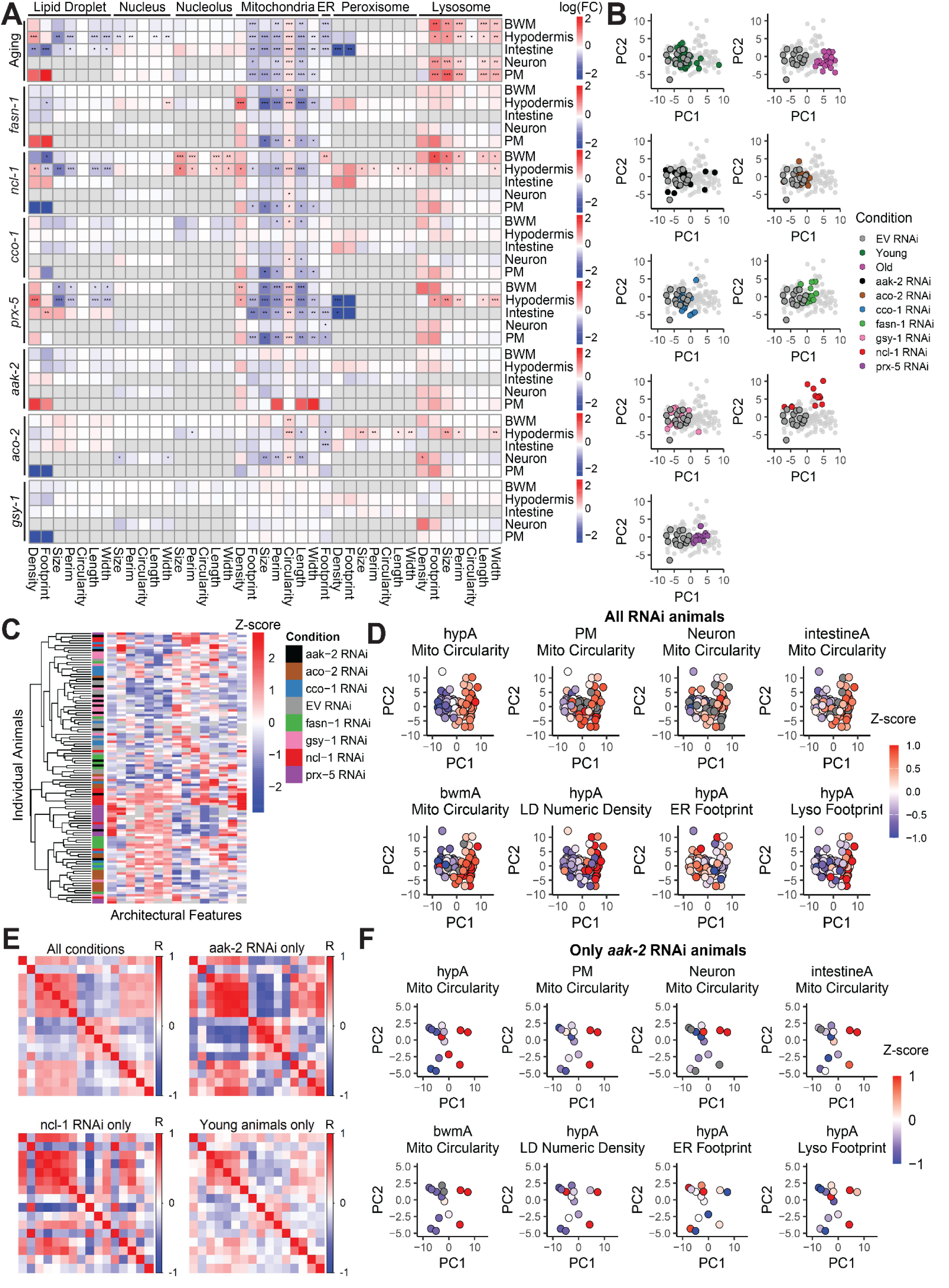
Many features of the MASS are common responses to disruption in homeostatic mechanisms and are frequently coregulated within individuals. A) Heat map showing the effect (log(FC)) of age (log(day12/day2)) or listed RNAi treatment (log(X RNAi/EV RNAi)) on subcellular architecture in MoMA *C. elegans* (n=166 animals, 13-27 animals per group, moderated t-test). B) Plot showing PC1 and PC2 from a PCA of all architectural endpoints analyzed across neuronal, PM, BWM, hypodermal and intestinal cells in MoMA *C. elegans* analyzing the effect of RNAi treatments and aging. Each plot shows EV RNAi animals (grey) relative to single conditions labeled in other designated colors (n=166 animals, 13-27 animals per group, PCA). C) Heatmap showing features of subcellular architecture which significantly change during aging (down columns) across individual animals (across rows), across MoMA *C. elegans* analyzed as a part of the RNAi screen (n=113 animals, 13-15 animals per group). D) Plot of PC1 and PC2 of a PCA analyzing all features of subcellular architecture analyzed across all RNAi treated animals colored by features of subcellular architecture. Each individual dot represents one individual animal (n=113 animals, 13-15 animals per group, PCA). E) Correlation matrices analyzing the correlation between features of subcellular architecture that change significantly during aging across either day 2, day 12 and all RNAi treatments combined (All conditions) or the designated subsets of animals listed in each respective plot title (n=113 animals, 13-15 animals per group, Pearson Correlation). F) Plot of PC1 and PC2 of the PCA shown in Fig. 4D, showing just *aak-2* RNAi treated animals, colored by features of subcellular architecture (n=14 animals, PCA). ∗ p < 0.05, ∗∗ p < 0.01, ∗∗∗ p < 0.001, for heatmaps, significance is capped at three asterisks.

We first asked how subcellular architecture changed across the RNAi treatments we analyzed and whether any perturbations elicited remodeling reminiscent of aging. Generally, we observed a broad range of effects: some RNAi clones produced minimal architectural changes (for example *aak-2* and *gsy-1* RNAi), whereas others drove widespread remodeling across multiple organelles and cell types (Fig. 4A). Although no single perturbation fully recapitulated the complete aging phenotype, several challenges convergently induced many MASS-like features. For instance, though effect size and significance was variable, mitochondrial fragmentation, reduced ER footprint and increased lysosome footprint were frequently observed across multiple distinct perturbations, such as *aco-2*, *prx-5*, and *ncl-1* RNAi (Fig. 4A). These data suggest that many features of subcellular architecture that change during the MASS represent, at least to some degree, generalized reactions to disruption in homeostatic mechanisms which can be driven by a diverse range of stimuli. However, other features, such as the decline in functional peroxisome abundance observed in intestinal cells, were not recapitulated across any challenges analyzed. Thus, some features of the MASS are somehow more specific to aging.

Fascinatingly, similar to the MASS itself, we found that many MASS features frequently developed together. At the population level, perturbations that induced one MASS-associated change often also induced many others (Fig. 4A). For example, conditions that promoted mitochondrial fragmentation in one cell type often also exhibited mitochondrial fragmentation across other cell types, together with lysosome expansion, reduced ER footprint, and increased hypodermal lipid droplet density. These relationships became even more apparent when analyzing subcellular architecture at the level of individual animals (Fig. 4B–E). Across all animals in all conditions analyzed, mitochondrial fragmentation in hypodermal cells was strongly correlated not only with mitochondrial fragmentation throughout the body but also with numerous architectural features of other organelles, indicating that remodeling extends across both organelle types and tissues (Fig. 4D, S4A). These relationships persisted even within genetically and environmentally identical populations. In conditions where animals appeared relatively heterogeneous, such as *aak-2* RNAi and day 2 animals, individuals exhibiting one MASS-like feature were likely to exhibit many others (Fig. 4F, S4B-D). Although we observed many exceptions, indicating that individual organelles retain some level of autonomy in their action, these data reveal a dominant pattern of coordinated remodeling across the body. Thus, our data collectively suggest that the MASS, at least in part, reflects the engagement of a generalized response to disruption in homeostatic mechanisms, which often manifests as a coordinated state switch in which many features are coregulated across the body, which could potentially be driven by a range of upstream events.

### The MASS resembles a reaction to physiological changes that develop in midlife during aging

We next were curious to understand what event(s) in midlife may cause the MASS. Interestingly, the window of time in which we observe the MASS is similar to a time in the *C. elegans* life cycle when they undergo many biological and physiological changes, such as a decline in locomotion, reduction in egg laying and a decrease in food consumption^30,42–47^. We were therefore next interested in determining whether any of these changes could be responsible for triggering the MASS. Toward this end, we first sought to establish the relationship between these physiological changes and the MASS. We tracked age-dependent physiological changes in MoMA animals across heterogeneous populations of day 9 adults, and then imaged subcellular architecture, attempting to link architectural features to the corresponding physiological measurements (Fig. 5A).

**Figure 5.**
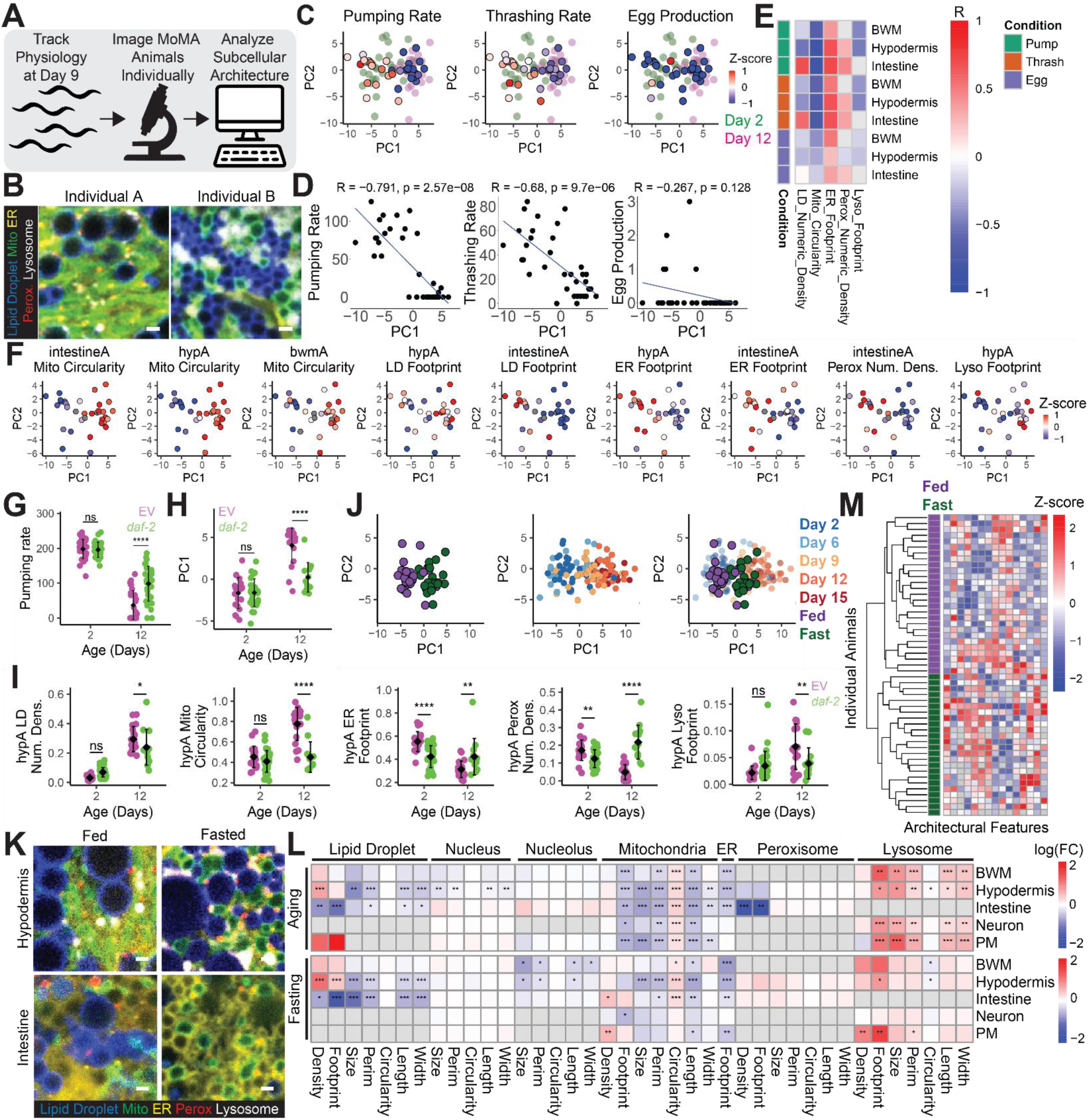
The MASS resembles a coordinated reaction to late life physiological state. A) Schematic outlining the experimental design to analyze correlation between subcellular architecture and animal physiology during aging. B) Images of anterior hypodermal cells of two distinct day 9 MoMA *C. elegans*. C) Plot showing PC1 and PC2 from a PCA of all architectural endpoints resolved across BWM, intestinal and hypodermal cells in day 2 and day 12 *C. elegans* (background; pink and green) together with Day 9 animals imaged with physiological parameters tracked (foreground; red-blue color scale) with thrashing, pharyngeal pumping rate, or egg laying measured in Day 9 animals colored by z score (n=88 animals, 26-35 animals per condition, PCA). D) Plots of thrashing, pharyngeal pumping rate or egg laying and PC1 from the PCA presented in Fig. 5C (n=35 animals, Pearson Correlation). E) Heat map plotting the correlation (R) between physiological measurements and architectural measurements across day 9 MoMA *C. elegans* (n=35 animals, Pearson Correlation). F) Plots of PC1 vs PC2 of a PCAanalyzing all day 9 animals analyzed shown in Fig. 5C colored by features of subcellular architecture (n=35 animals, PCA). G-I) MoMA *C. elegans* treated either with empty vector (EV) or *daf-2* RNAi were analyzed for pharyngeal pumping rate and subcellular architecture at day 2 and day 12 of adulthood (n=81 animals, 13-29 animals per condition, Two-way ANOVA with post-hoc Tukey’s test (G and I) or PCA(H), mean ± SD). J) Plot of PC1 and PC2 of a PCA analyzing all fed and fasted animals paired with animals across ages shown in Fig. 3 labeled either by age of animal or nutritional status (n=179 animals, 20-30 animals per condition, PCA). K) Images of anterior hypodermal and intestinal cells in MoMA *C. elegans* which were fed or fasted for 48 hours. L) Heat map showing log(FC) of architectural endpoints analyzed across day 2 and day 12 (log(day12/day2)) or across fed and 48 hour fasted (log(fasted/fed)) MoMA *C. elegans* anterior BWM, hypodermal, intestinal, neuronal and PM cells (n=104 animals, 24-27 animals per condition, moderated t-test). M) Heatmap showing features of subcellular architecture which significantly change during aging (down columns) across individual animals (across rows), with age annotated across fed or 48 hour fasted *C. elegans* (n=51 animals, 24-27 animals per condition). Scale bars, 1 µm. ∗ p < 0.05, ∗∗ p < 0.01, ∗∗∗ p < 0.001, ∗∗∗∗ p < 0.0001, for heatmaps, significance is capped at three asterisks.

Consistent with our earlier observations, day 9 animals exhibited substantial heterogeneity in subcellular architecture, with some individuals more closely resembling the majority of young animals and others more similar to old animals (Fig. 5B, S5A). By this time point, animals were largely no longer laying eggs, indicating that the MASS occurs after the midlife collapse in reproduction. In contrast, pharyngeal pumping and thrashing rates remained highly heterogeneous among day 9 individuals. Projecting each animal’s subcellular architecture onto PC1 revealed a strong and significant correlation between this axis and both pharyngeal pumping and thrashing rate (Fig. 5C–F, S5C): animals with more “aged-like” architectural features showed reduced pumping and motility, whereas those retaining “youthful-like” architecture maintained higher physiological performance. Further strengthening this association, we found that *daf-2* RNAi treated animals, in which this physiological collapse is delayed also exhibited an attenuation of the MASS at Day 12 (Fig. 5G-I). Together these findings establish an association between the MASS and the decline in organismal function observed during aging.

Finally, we sought to characterize this relationship between MASS and age-dependent physiological changes. Our data clearly establish a link between the two, but it remains unclear whether the MASS is a causal driver of these physiological changes, a downstream consequence of them, or a parallel process, with both potentially arising from a common upstream event. Toward this end, we used MoMA animals to track how subcellular architecture may remodel in response to fasting, mimicking the age-dependent decline in food consumption which develops in midlife alongside the MASS (Fig. 5J-M). Strikingly, we found that fasted subcellular architecture appeared remarkably similar to aging animals. For example, fasting and aging both increased hypodermal lipid droplet content, decreased intestinal lipid droplet content and drove mitochondrial fragmentation, reduction in ER footprint and expansion of the lysosome pool across many cell types. Much like our previous analyses, though imperfect, we again noted that animals that harbored one of these such changes tended to harbor many, indicating that fasting may also induce a subcellular architecture state switch similar to the MASS (Fig. 5M, S5D). Thus, these data suggest that the midlife shift in nutritional state that coincides with the MASS could be a driving force responsible for triggering MASS progression. Pairing these observations with our earlier data, these observations more broadly suggest that the MASS may constitute the engagement of a generalized structural state which forms as an adaptation to late life physiological challenges.

In sum, here we establish MoMA as an organism-level platform for mapping subcellular architecture across the body of a living animal. Though this architecture is highly dynamic in many respects, our data suggest that adult life is spent predominantly in one of two semi-discrete structural states, with the earlier portion of life spent in a youthful configuration, characterized by features such as elongated mitochondria, a small lysosome pool and expanded ER networks, and the later portion in an aged configuration, with fragmented mitochondria, an expanded lysosome pool and contracted ER networks. The transition between them, which we term the MASS, occurs relatively abruptly in midlife, and we show that it resembles a generalized response to a wide range of homeostatic challenges and can be triggered by physiological changes that develop alongside it. Our findings therefore raise the prospect that much of the age-dependent remodeling of subcellular architecture is not the outcome of stochastic local events accumulating over life within individual organelles and cells, but rather a coordinated, body-wide adaptation to late life physiological state. Moving forward, we anticipate that MoMA, the data we provide here and this conceptual framework will be valuable for continued efforts to decode both the biology of aging and the spatial regulation of cell biology.

## Discussion

Here we report the creation of MoMA, an organism-scale model of subcellular architecture which provides a systems-level view of how the structure of the cytoplasm is dynamically regulated across the body of fully intact, living animals. MoMA is a single *C. elegans* strain that stably expresses all transgenes used to visualize subcellular architecture and can be imaged on several common types of fluorescent microscopes simply following animal immobilization, without additional time-or cost-intensive sample processing, making MoMA a highly accessible system. A central advantage of MoMA is that every measurement remains indexed to the organelle, cell type, body region and individual animal it came from, so the behavior of any one organelle can be analyzed against the backdrop of every other organelle measured elsewhere in that same animal. Though our initial focus has been on organelle abundance and morphology in the context of aging, MoMA can be applied to explore many other dimensions of subcellular architecture, such as organelle-organelle interactions and organelle motility, across virtually any physiological condition. Because MoMA scales this measurement across whole animals rather than single cells, the datasets it produces are well suited for training artificial intelligence–based, organism-level models of subcellular structure, and ultimately for predictive, *in silico* “virtual cells” that forecast how cytoplasmic organization responds to physiological change. As such, we expect MoMA and the data we provide in this manuscript to be a broadly valuable asset in ongoing efforts to decode the spatial logic of cell biology. Here we discuss how we initially employed MoMA to uncover principles underlying how and why subcellular architecture is dynamically regulated during aging and how this reshapes our general understanding of both aging and the spatial regulation of cell biology.

First, we provide a global view of how subcellular architecture changes during aging, allowing us to identify the extent to which changes develop, resolve how these changes unfold spatially and temporally throughout the body and, by extension, to refine our understanding of both why they arise and their functional implications. Although subcellular architecture is in many respects cell-type specific, we find that many age-induced changes are broadly conserved throughout many cell types and develop generally consistently across individuals. Moreover, these changes predominantly emerge in midlife at an inflection point, relatively synchronously within individuals, as part of an event we term the MASS. Because subcellular architecture is not perfectly synchronized as animals transition through the MASS (for instance, fragmented mitochondria in one cell do not always perfectly predict a contracted ER or expanded lysosome pool in that same cell), our data indicate that organelles remain predominantly governed by local mechanisms operating at the level of individual organelles within individual cells. Yet the pervasive correlation between features of subcellular architecture across the body within individuals suggests that these local mechanisms are themselves often coordinated by events experienced systemically throughout the body. Consistent with this, we find that the MASS correlates strongly with age-dependent changes in physiological function that impact the body systemically and we show that much of the MASS can be triggered by this change in physiological state. However, despite this within-individual coordination, the timing of the MASS varies across individuals. Collectively, these observations suggest that age-dependent changes in subcellular architecture may largely develop acutely in midlife as a reaction to changes in physiological state, induced by an upstream event that onsets stochastically across individuals (Fig. S5E). This model pairs well with prior work showing that many MASS features, such as induction of autophagy and mitochondrial fragmentation, constitute protective adaptations to a range of homeostatic challenges, such as heat shock and starvation^48–50^.

Our study also contributes to a broader ongoing effort to understand how the biology of aging unfolds across the body of an animal. Original ideas of aging favored the view that aging is a highly stochastic process in which biological systems progressively wear down over time. This idea is supported by many studies that resolve a progressive and increasingly stochastic drift in biological phenotypes during aging^30,31,51,52^. Though newer studies have not refuted this view, we now know that aging also involves more intricate patterns of change, for instance, phenotypes that develop at different rates across different parts of the body or that emerge non-linearly in relatively abrupt bursts^52–56^. Energy expenditure in humans, for example, remains relatively consistent through adult life before beginning to drop around age 60^57^. Moreover, menopause in females is linked to abrupt shifts in endocrine, skeletal, hepatic, renal and lipid systems^58^. Our identification of the MASS builds upon this work by showing that these non-linear waves of change are not necessarily restricted to subsets of tissues or cell types, but can in at least some cases reflect coordinated, organism-wide shifts in state. Further, our data suggest that these non-linear shifts in state could possibly result from physiological changes that develop acutely in midlife during aging. It is plausible that other features of cell biology change in similar ways to the MASS, and indeed many existing datasets are consistent with this same pattern of coordinated, non-linear emergence. For example, many aging clocks have recently been appreciated to resolve forms of stress which can develop and resolve transiently throughout life^59^. However, new tools that provide both organism-level coverage and preservation of spatial resolution, similar to MoMA, will need to be developed to empirically evaluate this possibility.

These results also evolve our general understanding of the spatial organization of the cytoplasm by revealing an unexpected single axis that captures a substantial fraction of variation in subcellular architecture throughout the body across diverse spatial and physiological contexts. Specifically, we found that many features of subcellular architecture that appear during aging are broadly inducible by a range of homeostatic challenges, and that these changes are frequently coregulated with one another. This coregulation is loosely apparent when analyzing animals as bulk populations. For example, mitochondrial fragmentation in one cell type under one challenge often coincides with mitochondrial fragmentation in other cell types and with other MASS-like changes, such as expansion of the lysosome pool and ER contraction. However, this pattern becomes much more striking when analyzing animals individually, as many features of the MASS are significantly correlated with one another across the body, across and within a wide range of homeostatic challenges. Thus, although our data clearly indicate that individual organelles preserve some level of autonomy in their action, they also suggest that *C. elegans* often loosely oscillate between two dominant architectural configurations. Based on the conditions under which animals occupy each state and the combination of features that characterize them, we speculate that these states may be optimized either for growth, reproduction, and anabolism (a youthful, fed, or control-treated subcellular architecture – characterized by features such as elongated mitochondria, expanded ER networks and low levels of lysosomes) or for survival and catabolism (a aged, fasted, or challenged subcellular architecture – characterized by features such as fragmented mitochondria, a contracted ER network and an expansion of the lysosome pool). Applied to aging, this framework suggests that aging involves a switch between two states. Young animals occupy the growth/reproduction state basally, but are primed to dynamically switch to the survival state when needed. In response to challenges that arise during aging there is a rapid shift to the survival state in midlife. In this model, the MASS is a protective response to aging, but one that leaves the cell less able to dynamically response to further challenges.

Together, as these discoveries were only possible by leveraging the unique combination of technical strengths MoMA provides, these findings illustrate the power of MoMA to reveal fundamental principles governing the spatial regulation of cell biology, not only during aging but across many aspects of health and disease more broadly. As MoMA is applied to analyze new physiological contexts and additional dimensions of subcellular architecture, we anticipate it will continue to reveal general organizing principles of cellular spatial organization which can transform our understanding of cell biology, organismal physiology and disease.

## Contributions

C.S.M. and W.B.M. conceived the idea for the study and C.S.M. conducted and analyzed all experiments not listed hereafter. A.P. and J.D.K. assisted with assay validation and statistical analyses. C.G.S.G. constructed the lipid droplet reporter. P.V.A. and P.M.L. assisted with microscopy. J.M. assisted with statistical analyses. W.B.M. oversaw and conceptually developed the study. C.S.M. wrote the manuscript with input from all authors.

## Acknowledgements

CSM is funded by F32AG085703 and K99AG092471 from NIH/NIA. W.B.M. is funded by R01AG067106 and R01AG044346 from NIH/NIA. We thank Maxwell G. Heiman and Adrià Razzauti for help with strain integration. We thank all members of the Mair laboratory for their support.

## Materials and Methods

### C. elegans strains and husbandry

Strains used in this study were N2 (WT); WBM1472 *plin-1(wbm78[wrmBFP::plin-1]) I*; WBM1882 *N2, wbmEx455[eft-3p::his-24::eCFP]*; WBM1231 *N2, wbmIs97[eft-3p::tomm20::GFP::unc-54 3′UTR, wbmIs65]*; WBM1875 *N2, wbmEx454[eft-3p::TRAP-1::hfYFP]*; WBM1873 *N2, wbmEx456[eft-3p::CemOrange2::SKL]*; WBM1874 *N2, wbmEx452[eft-3p::NUC-1::mCherry]*; WBM1880 *N2, plin-1(wbm78[wrmBFP::plin-1]) I, wbmIs97[eft-3p::tomm20::GFP::unc-54 3′UTR, wbmIs65], wbmIs168[HIS-24::CFP, TRAP-1::hfYFP, CemOrange2::SKL, NUC-1::mCherry]*; WBM1871 *N2, plin-1(wbm78[wrmBFP::plin-1]) I, wbmIs97[eft-3p::tomm20::GFP::unc-54 3′UTR, wbmIs65], wbmIs168[HIS-24::CFP, TRAP-1::hfYFP, CemOrange2::SKL, NUC-1::mCherry]*; ABR161 *hjIs37[vha-6p::mRFP-PTS1 + Cbr-unc-119(+)]*; *ldrIs1[dhs-3p::dhs-3::GFP + unc-76(+)]*; IK374 *tax-6(p675)*; *njEx220[odr-3p::tax-6 cDNA, myo-3p::tax-6 cDNA, myo-3p::GFP, ges-1p::NLS-GFP]*; WBM1688 *scpl-4(wbm108[TIMM50::scarlet]) V*; VK2748 *vkEx2748[nhx-2p::CemOrange2::tram-1 + nhx-2p::GFP::KDEL]*; VS11 *hjIs73[vha-6p::GFP::daf-22 + C. briggsae unc-119(+)]*; WBM1215 *N2, wbmIs89[rab-3p::3XFLAG::dpy-10::SL2::wrmScarlet::rab-3 3′UTR, wbmIs68] IV*; GRU101 *gnaIs1[myo-2p::YFP]*; WBM1633 *N2, wbmIs153[dpy-7p::3XFLAG::dpy-10::SL2::wrmScarlet::unc-54 3′UTR, wbmIs88] V*; WBM1133 *N2, wbmIs63[myo-3p::3XFLAG::wrmScarlet::unc-54 3′UTR, wbmIs61]*; WBM1478 *N2, wbmIs136[nep- 17p(mini)::3XFLAG::dpy-10::SL2::wrmScarlet::unc-54 3′UTR, wbmIs88] V*; COP262 *unc-119(ed3) III*; *knuSi221[fib-1p::fib-1(genomic)::eGFP::fib-1 3′UTR + unc-119(+)]*; WBM1795 *N2, plin-1(wbm78[wrmBFP::plin-1]) I, wbmIs97[eft-3p::tomm20::GFP::unc-54 3’UTR, wbmIs65]*; WBM1860 *N2, plin-1(wbm78[wrmBFP::plin-1]) I, wbmIs97[eft-3p::tomm20::GFP::unc-54 3’UTR, wbmIs65], wbmEx448[HIS-24:CFP, TRAP-1:hfYFP, CemOrange2:SKL, NUC-1:mCherry]*. *C. elegans* were maintained on standard nematode growth media (NGM) seeded with *E. coli* (OP50-1) at 20 °C. OP50-1 was cultured overnight in LB at 37 °C and then added to NGM plates and allowed to grow for 2 days prior to adding *C. elegans* to each plate. To generate day 2 adults, a timed egg lay was performed and L4 larvae were picked 48 h prior to imaging. Aged animal cohorts were transferred at least once every other day to avoid starvation.

### *C. elegans* Perturbation Experiments

Animals were treated with RNAi by feeding animals RNAi from hatch via supplementing animal food with plasmids engineered to express each RNAi clone respectively (HT115 bacteria in the L4440 vector). RNAi clones were obtained either from the Rual et al 2004 or Kamath et al 2003 libraries^60,61^. As previously described^62,63^, prior to moving animals to plates 0.1 M isopropylthiogalactoside (IPTG) was added to each plate for a final concentration in each plate of 1 mM IPTG^64^. To fast *C. elegans*, animals were raised until day 1 of adulthood and then transitioned to plates with NGM plates seeded with OP50-1 and containing 40 µM 5-fluoro-2′-deoxyuridine (FUDR) for 24 hours. At day 2 of adulthood, animals were then transferred to NGM plates with 40 µM FUDR and no OP50-1 and transferred at least once every 24 hours up until imaging. Fasted animals were then compared against fed control animals, always exposed to food, which were also treated identically with FUDR.

### DNA Cloning

Plasmids created in this study were: pCM26 (*eft-3p::CemOrange2::SKL*); pCM27 (*eft-3p::nuc-1::mCherry*); pCM31 (*eft-3p::his-24::CFP*); and pCM37 (*eft-3p::trap-1::hfYFP*). To create pCM26, *CemOrange2*^65^ was amplified using AGTTGGAATTCTTAcaacttggaCTTGTAGAGTTCGTCCATTCCTCC and TGAGGGTACCATGGTCTCCAAGGGAGAGGAGAAC. The resulting amplicon and pSD1 were then digested with KpnI and EcoRI and ligated together. To create pCM27, *nuc-1* was amplified out of WT *C. elegans* cDNA first using a primary PCR with ATGGGCTTGTCTCCTGCCGCTGTGC and TGCACAATTATTTTGGGTTGCAATTACACTGTAGGCTTTCCAAAGGA followed by a second PCR with CTCACCCGGGATGGGCTTGTCTCCTGC and ccatCGGTACCCCTGCACAATTATTTTGGGTTGCAATTACA. The resulting amplicon and pCH2 were then digested with KpnI and XmaI and ligated together. To create pCM31 *his-24* was amplified out of WT *C. elegans* cDNA first using a primary PCR with ATGTCTGATTCCGCTGTTGTTGCCGCCG and GGCCTTGGCGGCTGGCTTGGC followed by a second PCR with CTCACCCGGGATGTCTGATTCCGCTGTTGTTG and CCATGGTACCGGCCTTGGCGGCTG. The resulting amplicon and pCM22 were then digested with XmaI and KpnI and ligated together. To create pCM37 *trap-1* was amplified out of WT *C. elegans* cDNA using CTTGAGGGTACCATGAAGCTGTCGACCGTCTTTCTTCTC and ttgcgaccggtACGTCAGCCTTCTTGGC. The resulting amplicon and pCM29 were then digested with KpnI and AgeI and ligated together. To create a homology directed repair (HDR) template for creating WBM1472, *wrmBFP* was amplified from pUA97 using gaacgtattttctagtttatcattttatcaattgattgtttgcagcaagaaagATGTCAGAGCTTATTAAGGAGAATATG and CTAAGACTTGATCGTAGTAGCTTTGGGCTTGGGCTTGTTGATCCTCAACcGATACaGGC TGtTCGACGTCAGTATTAAGCTTGTGACCCAGTTTGC.

### *C. elegans* Strain Creation

Following established protocols^66^, homology directed repair templates were generated by PCR with primers that attach at least 35 base pairs of homology on either end of the insert. CRISPR injection mixes were generated with the following composition: 0.375 μl Hepes pH 7.4 (200 mM), 0.25 μl KCl (1 M), 2.5 μl tracrRNA (4 μg/μl), 0.6 μl crRNA (2.6 μg/μl), HDR template 500 ng/μl, water up to 8 μl. Prior to injection, 2 μl purified Cas9 (12 μg/μl) was added before the solution was centrifuged at 13,000 rotations per minute (rpm) and then incubated at 37 °C for 10 min. *C. elegans* extrachromosomal array mutants were generated by injecting 30 ng/µl plasmid DNA in water purified using Zymo D4019. To create WBM1860 (non-integrated MoMA), WBM1795 N2, *plin-1(wbm78[wrmBFP::plin-1]) I, wbmIs97[eft-3p::tomm20::GFP::unc-54 3’UTR, *wbmIs65]* was injected with pCM26, pCM27, pCM31 and pCM37. To integrate WBM1860 for more stabilized expression of MoMA reporters across the body and produce WBM1880, WBM1860 *C. elegans* were incubated with 10 µg/ml psoralen for 15 minutes in the dark at room temperature, irradiated at 360 µJules x 100 and then screened for successful integrants. All newly generated strains were outcrossed at least 6 times prior to using in experiments.

### Microscopy

For imaging, *C. elegans* were placed in 0.1 µm PolyBead Microspheres (PolyScience 00876-15) on a 10% agar pad, covered with a #1.5 coverslip and imaged immediately after mounting. All images were acquired using an upright Leica Stellaris 8 DIVE FALCON 2-photon + confocal microscope outfitted with 4Tune spectral NDD and an InSight X3 dual beam pulsed (80 MHz) infrared laser (Spectra Physics) (Fig. S1-C) and a pulsed WLL (80 MHz) or a Zeiss LSM980 outfitted with 2x multi-alkali PMT + GaAsP 32 channel spectral detector and 405, 458, 488, 514, 561, 594, 639 and 730 nm laser lines (All other figures, except for Fig. S1-C). On the Leica Stellaris, images were acquired with a Plan Apo 63x oil-immersion objective with 1.4 numerical aperture at 2x zoom, 512 × 512 (effective pixel size: 172 nm) pixel images and a pixel dwell of 3.16 µs. LasX acquisition software was used to control the microscope. wrmBFP was imaged using 405 nm Ex, 410-450 nm Em; eCFP was imaged using 441 nm Ex, 452-480 nm Em; GFP was imaged using 484 nm Ex, 488-505 nm Em; hfYFP was imaged using 514 nm Ex, 520-560 nm Em; CemOrange2 was imaged using 530 nm Ex, 540-560 nm Em; mCherry was imaged using 605 nm Ex, 625-728 nm Em. On the Zeiss LSM980, images were acquired with a 63x oil-immersion objective with 1.4 numerical aperture at 2x zoom, 954 x 954 (effective pixel size ∼70 nm) and unidirectional scanning at scan speed 6. Zen was used to control the microscope. Images were collected by exciting with 405 nm, 445 nm, 514 nm, 561 and 639 nm lasers simultaneously and collecting emission across ∼410-694 nm in ∼10 nm bins in the GAaSP 32 channel spectral detector. Linear unmixing was performed in Zen. In this study, all analyses were performed on two-dimensional single z-slices to capture a representative section of the cytoplasm. For all imaging experiments each condition was replicated at least 3 times.

### Image Analysis

After unmixing, images were analyzed in ImageJ/FIJI using a semiautomated analysis pipeline designed to quantify morphologic features of organelles and then data were further processed in R. To analyze subcellular architecture in ImageJ/FIJI, first, a region of interest was manually drawn to isolate a specific cell type to be analyzed within the *C. elegans* body, then organelles were analyzed sequentially using the following protocols. For all organelle measurements, Set Measurements was configured to: run(“Set Measurements…”, “area perimeter bounding shape feret’s integrated display redirect=None decimal=3”). To analyze peroxisomes, lysosomes and ER, a threshold was set to isolate each signal and then Analyze Particles was used with size=0.002-Infinity to quantify particle morphology. To analyze lipid droplets, nuclei, nucleoli and mitochondria, organelles were manually traced. The lysosome reporter in intestinal cells was unexpectedly most often not detectable. Therefore, we omitted analysis of intestinal lysosomes in this study. Lysosomal gut autofluorescent granules were detected in the lipid droplet and nuclear channels, but were easily discriminable from lipid droplets and nuclei based on cross channel fluorescence and morphological differences. Nucleoli were only apparent in a subpopulation of HIS-24::eCFP labeled nuclei. Therefore, some cells which have a nuclei measured do not have a corresponding nucleolar measurement. Peroxisomal reporter fluorescence persists after degradation of peroxisomes via pexophagy^6,67^. Thus, lysosomes are sometimes CemOrange2+/mCherry+ and peroxisomes were determined to be structures that were CemOrange2+/mCherry-. Cell type identification was validated using fluorescent reporters of each respective cell type: WBM1215 N2, *wbmIs89[rab-3p::3XFLAG::dpy-10::SL2::wrmScarlet::rab-3 3′UTR, wbmIs68] IV*; GRU101 *gnaIs1[myo-2p::YFP]*; WBM1633 N2, *wbmIs153[dpy-7p::3XFLAG::dpy-10::SL2::wrmScarlet::unc-54 3′UTR, wbmIs88] V*; WBM1133 N2, *wbmIs63[myo-3p::3XFLAG::wrmScarlet::unc-54 3′UTR, wbmIs61]*; WBM1478 N2, *wbmIs136[nep-17p(mini)::3XFLAG::dpy-10::SL2::wrmScarlet::unc-54 3′UTR, wbmIs88] V*. Body region is denoted in the figures in many listed endpoints either with an “A” to denote anterior or “P” to denote posterior (e.g. hypA refers to anterior hypodermis).

### Statistical analyses and reproducibility

All experiments were repeated at least three times. “n” denotes the number of animals or cells analyzed as indicated in each respective figure legend. All statistical tests used are indicated in the figure legend for each respective analysis. In general, to evaluate statistical significance in observed differences between two groups, Student’s t-test was used; to evaluate statistical significance in observed differences between more than two groups, a two-way ANOVA with post-hoc Tukey’s test was used. All analyses were performed either in GraphPad Prism or RStudio (2023.3.1.446) using R version 4.5.1 (2025-06-13). To compute organelle footprint, the area occupied by each organelle was divided by the total area of the region analyzed. To compute organelle density by number, the number of particles analyzed was divided by the total area of the region analyzed. Measurements per cell were made by averaging values across all organelles of a specific type in each cell respectively. PCAs were performed using prcomp() in R with missing values imputed when relevant. Heatmaps were made using pheatmap() (version 1.0.12) in R and show either the log(fold change (FC)), Z-score, or Pearson’s correlation coefficient (R). Differential features were identified using linear models with empirical Bayes–moderated t-statistics (limma; version 3.50.3), and significance was determined after Benjamini–Hochberg false discovery rate correction across all tested features. Pearson’s correlation analyses were performed using cor() in R. ChatGPT and Claude were used to help develop and edit code used in this manuscript.

### Lysotracker Labeling

To label lysosomes with Lysotracker Green, 50 µM Lysotracker Green in M9 was added to OP50-1 NGM plates and allowed to dry. Then animals were placed on plates with Lysotracker Green and incubated at 20°C overnight prior to imaging to achieve labeling.

### Lifespan and Healthspan Experiments

Experiments were conducted on NGM with 100 µl OP50-1 seeded 48 hr at room temperature before adding animals. *C. elegans* were synchronized by egg lay and ∼48 h later, L4-stage *C. elegans* were picked to have day 1 adults to prepare for each respective experiment. To analyze physiological properties, *C. elegans* were generated as described above, and then analyzed for pharyngeal pumping, egg laying and thrashing rate. To analyze thrashing rate, *C. elegans* were placed in M9 solution, acclimated for 30 seconds, and then monitored for thrashes per minute. To track pumping rate, *C. elegans* were scored for pumps per minute on bacterial lawns. To analyze egg laying, animals were plated on single plates for 24 hours and then scored for number of eggs laid per animal.

### Food Ingestion Assays

Animals were scored for pharyngeal pumping rate and placed into either a high pumping rate (greater than or equal to 24 pumps per minute) or low pumping rate (less than 24 pumps per minute group). Animals were then placed onto OP50-1 seeded plates pretreated by adding 100 µl 0.1 µm Fluoresbrite YG Microspheres (Polysciences 21636-1; diluted 1:5 in M9 buffer) on top of the bacterial lawn. Animals were allowed to eat for 5 minutes before anesthetization and imaging under a fluorescent stereoscope. To analyze data, a region of interest was drawn around the pharyngeal lumen to measure amount of ingested food. Ingested fluorescent food was then analyzed with each region of interest normalized to area with background fluorescence subtracted.

### LLM Use

ChatGPT and Claude were used to edit the manuscript.

---

**Figure S1.**
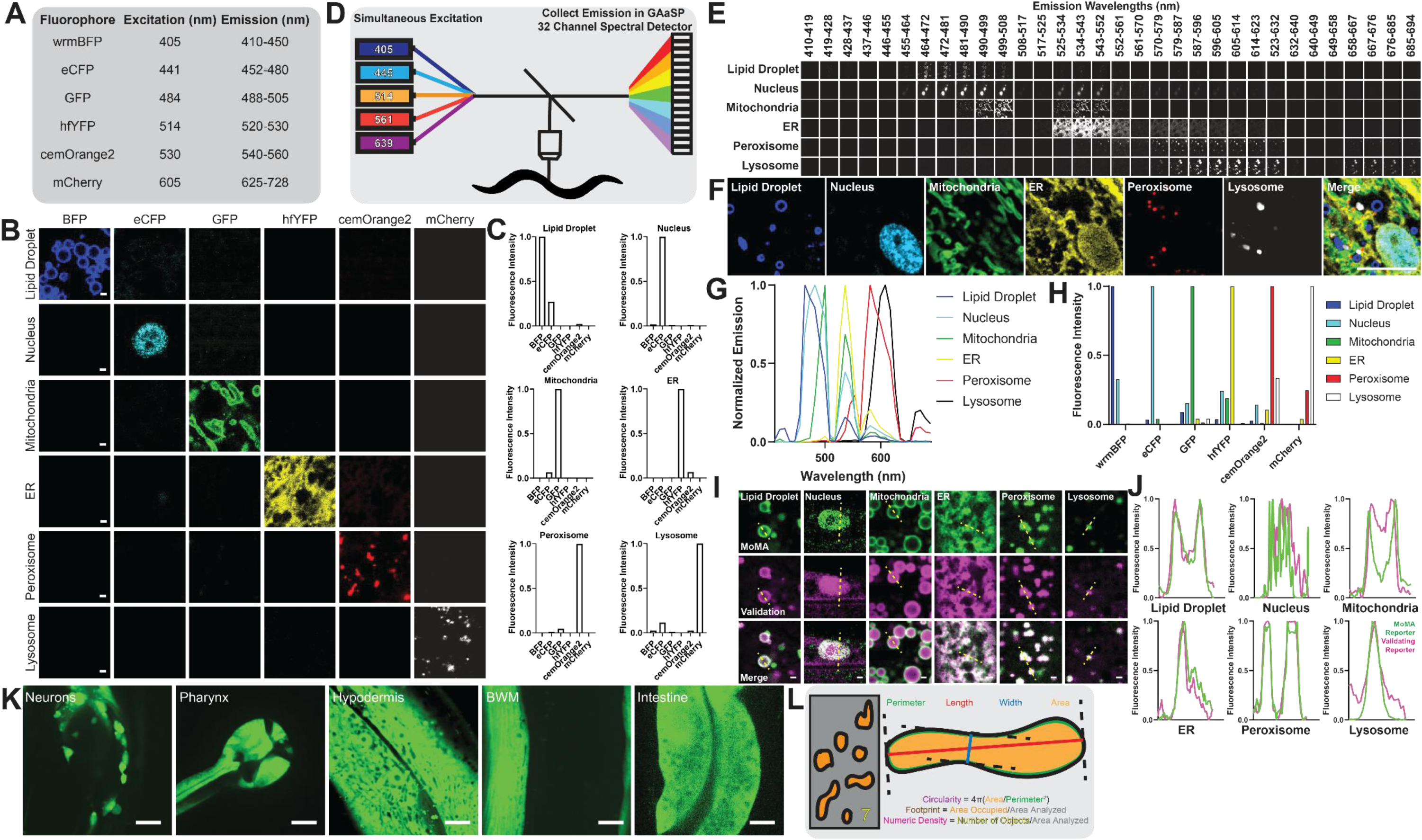
Confocal Imaging Strategy 1: A) Table listing excitation and emission wavelengths used to distinguish fluorescent proteins in MoMA. B-C) Images and quantification of day 2 adult *C. elegans* strains including each MoMA organelle reporter individually imaged using the specifications listed in Fig. S1A: (wrmBFP::PLIN-1; labeling lipid droplets), HIS-24::eCFP; labeling nuclei), TOMM-20_aa1-49_::GFP; labeling mitochondria), TRAP-1::hfYFP; labeling the endoplasmic reticulum), CemOrange2::SKL; labeling peroxisomes) and NUC-1::mCherry, labeling lysosomes). Confocal Imaging Strategy 2: D) Schematic of spectral imaging strategy to image MoMA. E) Images of day 2 adult *C. elegans* including each MoMA organelle reporter individually imaged using the specifications listed in Fig. S1D: (wrmBFP::PLIN-1; labeling lipid droplets), HIS-24::eCFP; labeling nuclei), TOMM-20_aa1-49_::GFP; labeling mitochondria), TRAP-1::hfYFP; labeling the endoplasmic reticulum), CemOrange2::SKL; labeling peroxisomes) and NUC-1::mCherry, labeling lysosomes). F) Image of day 2 adult hypodermal MoMA *C. elegans* cell showing lipid droplets, nuclei, mitochondria, ER, peroxisome and lysosomes unmixed through linear unmixing. G) Fluorescence emission spectra for all MoMA probes. H) Fluorescence intensity of each MoMA probe across all unmixed channels. I-J) Images and quantification of fluorescence intensity of MoMA reporters imaged with alternative markers of each organelle imaged in MoMA. Alternative markers imaged include: 1) lipid droplets, DHS-3::GFP, intestine, 2) nuclei, NLS::GFP, intestine, 3) mitochondria, SCPL-4::wrmScarlet, BWM, 4) ER, CemOrange2::TRAM 1, intestine, 5) peroxisome, GFP::daf-22, intestine, 6) lysosome, Lysotracker Green, hypodermis. K) Images of day 2 adult *C. elegans* harboring cassettes for tissue specific expression of a fluorescent protein localized to the cytosol of neuronal, intestinal, hypodermal, PM or BWM cells. L) Schematic outlining endpoints of geometric analysis of organelle architecture. Scale bars, 1 µm (B, I), 10 µm (F, K).

**Figure S2.**
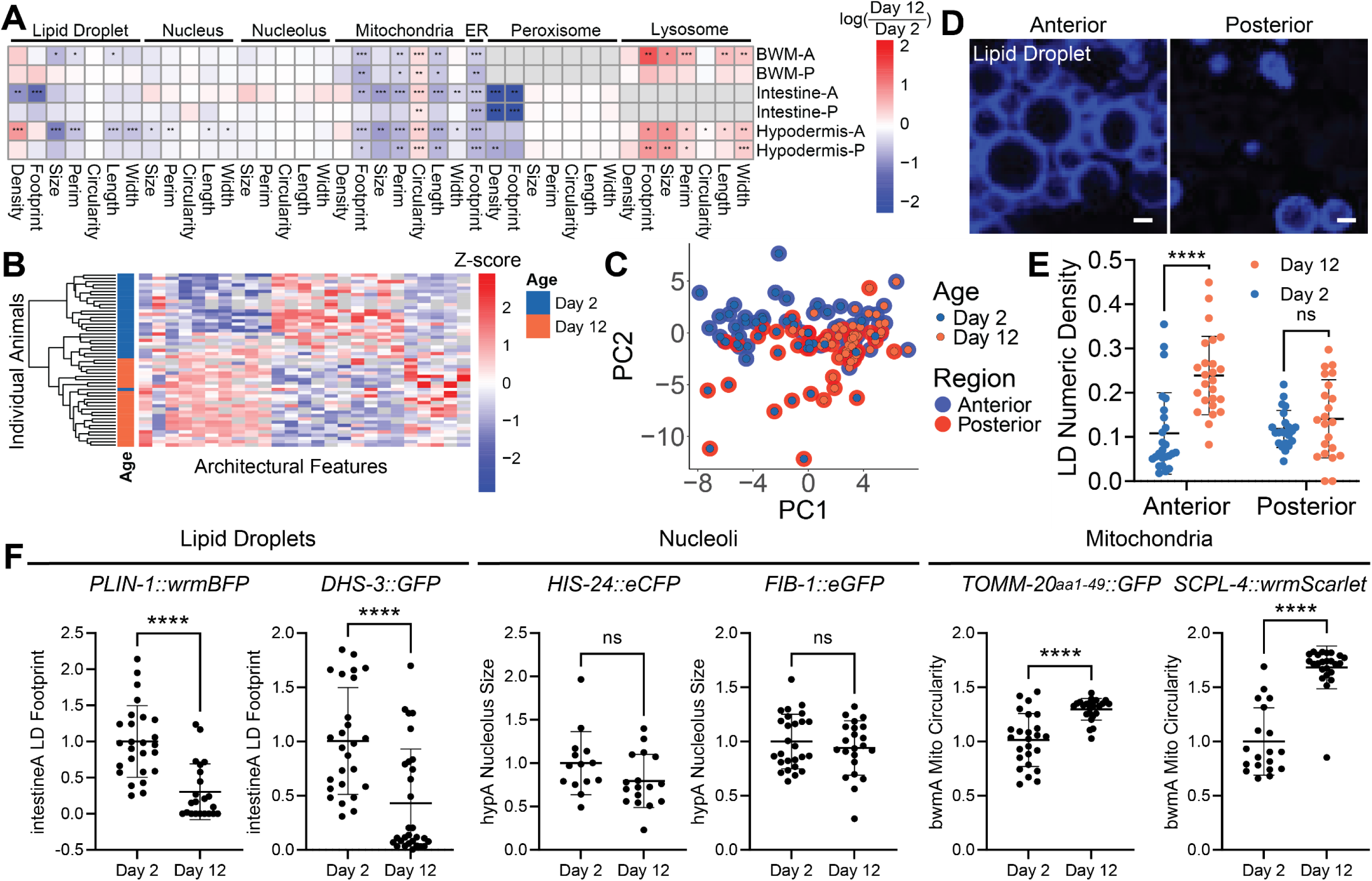
A) Heat map showing the log(day 12/day 2) of architectural endpoints across day 2 and day 12 animals analyzed in MoMA *C. elegans* across anterior BWM, hypodermal, intestinal anterior and posterior cells (n=53 animals, 26-27 animals per age, moderated t-test). B) Heatmap showing features of subcellular architecture which significantly change during aging (down columns) across individual animals (across rows), with age annotated (n=53 animals, 26-27 animals per age). C) PCA analyzing animal body regions of day 2 and day 12 animals reflecting all features of subcellular architecture analyzed per body region in BWM, intestinal and hypodermal cells (n=53 animals, 26-27 animals per age, PCA). D-E) Images and plot showing LD density in day 12 (and day 2 in E) MoMA *C. elegans* across either anterior or posterior cells (n=22-25 animals per group, Two-way ANOVA with post-hoc Tukey’s test; mean ± SD). F) Plots showing analysis of architectural endpoints of MoMA *C. elegans* compared against alternative labels for each organelle respectively across young (day 2) or old (day 12) *C. elegans*. Reporters used were: lipid droplets, DHS-3::GFP, wrmBFP::PLIN-1; nucleoli, FIB-1::eGFP, HIS-24::eCFP; mitochondria, SCPL-4::wrmScarlet, TOMM-20_aa1-49_::GFP (n=14-26 animals per condition, Student’s t-test, mean ± SD). ∗ p < 0.05, ∗∗ p < 0.01, ∗∗∗ p < 0.001, ∗∗∗∗ p < 0.0001, for heatmaps, significance is capped at three asterisks.

**Figure S3.**
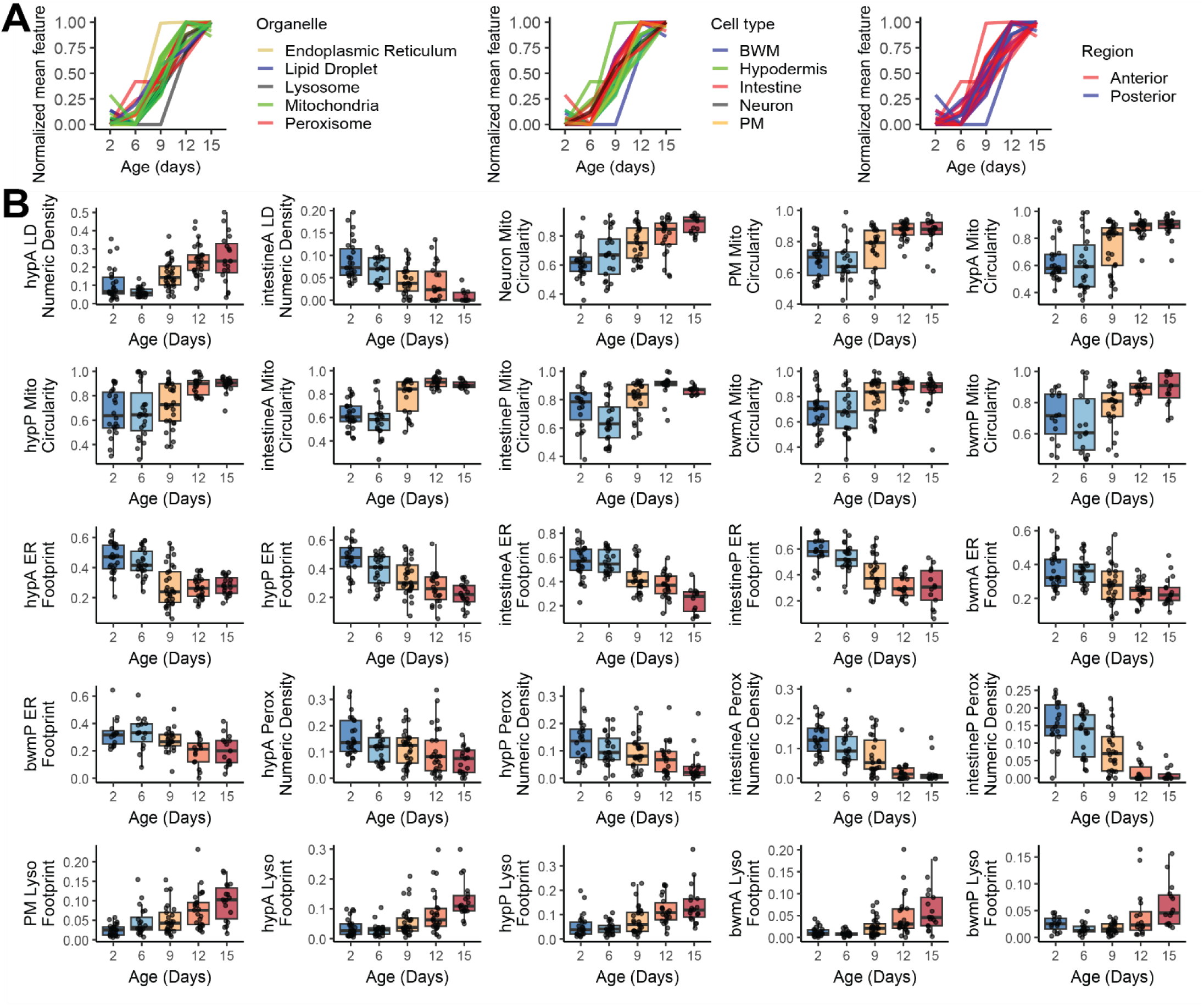
A) Plots of features of subcellular architecture which significantly change during aging across day 2, 6, 9, 12 and 15 animals with youthful values set to 0 and aged values set to 1 and lines colored by either organelle, cell type or body region (n=128 animals, 20-30 animals per age). B) Plots of individual features of subcellular architecture across day 2, 6, 9, 12 and 15 MoMA *C. elegans* (n=128 animals, 20-30 animals per age, box shows median and interquartile range).

**Figure S4.**
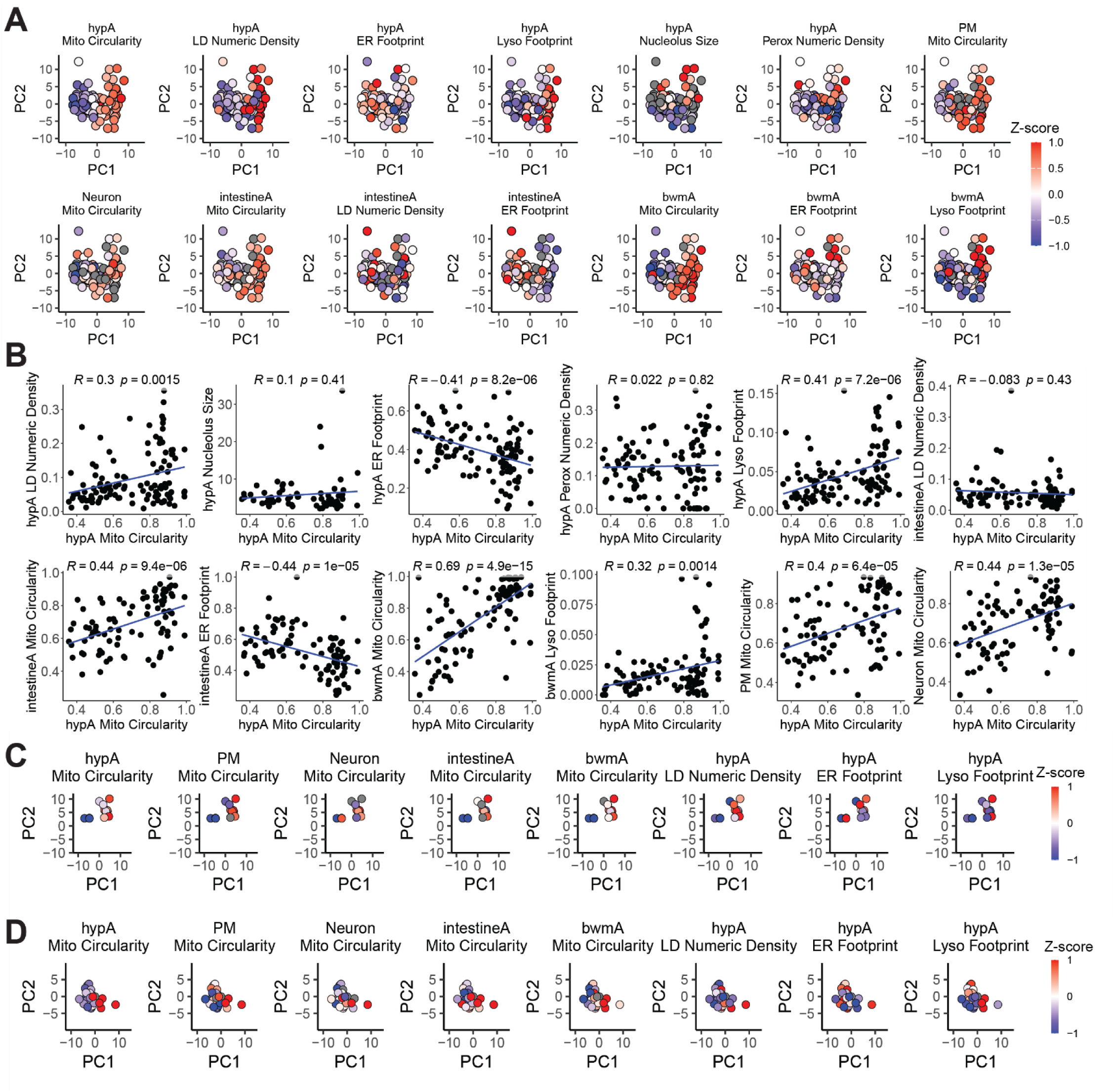
A) Plot of PC1 and PC2 of a PCA analyzing all features of subcellular architecture analyzed across all RNAi treated animals colored by features of subcellular architecture. Each individual dot represents one individual animal (n=113 animals, 13-15 animals per group, PCA). B) Correlation plots comparing anterior hypodermal mitochondrial circularity to other features of subcellular architecture across the body of animals across all RNAi treated animals (n=113 animals). C-D) Plot of PC1 and PC2 of the PCA shown in Fig. 4D, showing just *ncl-1* RNAi treated (C) or day 2 (D) animals, colored by features of subcellular architecture (n=13 (C) or 26 (D), PCA).

**Figure S5.**
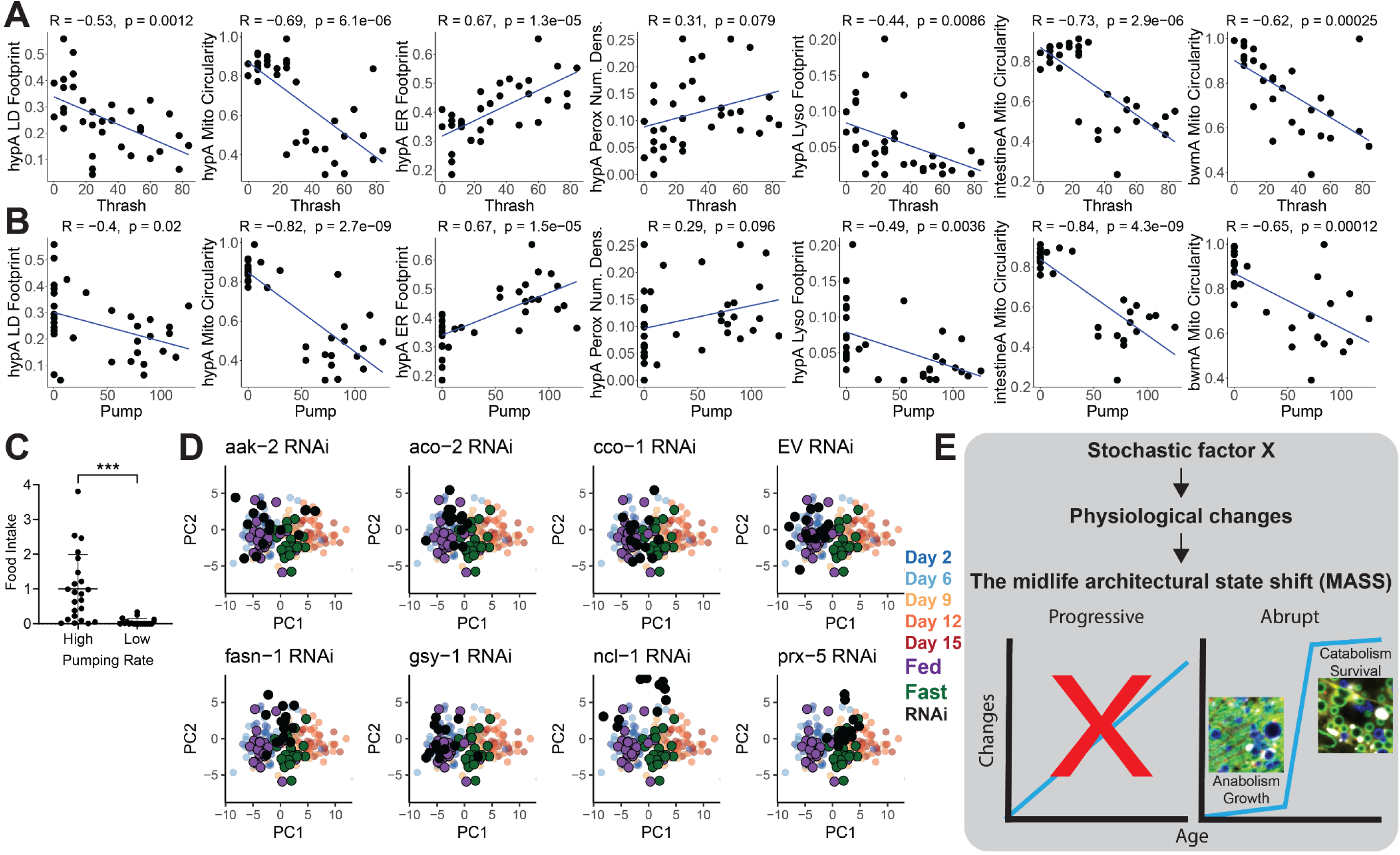
A-B) Plots of thrashing (A) or pharyngeal pumping rate (B) compared against features of subcellular architecture in day 9 MoMA *C. elegans*. Each dot represents an individual animal (n=35 animals, Pearson Correlation). C) Analysis of food consumption, tracked by ingestion of fluorescent bead labeled OP50-1 in Day 9 *C. elegans* split into groups by either high or low pumping rate (n=41 animals, 18-23 animals per condition, Student’s t-test, mean ± SD). D) Plot of PC1 and PC2 of a PCA analyzing all animals of all ages and RNAi treatments shown in Fig. 3 and 4 compared to the fed and fasted animals analyzed in Fig. 5 (n=292 animals, 13-30 animals per condition, PCA). E) Cartoon outlining a working model for the cause and development of the MASS. ∗∗∗ p < 0.001, ∗∗∗∗ p < 0.0001.

